# Modelling the formation of skin ulcers observed in Buruli ulcer disease

**DOI:** 10.64898/2026.10.06.757134

**Authors:** Fatumah Atuhaire, Rachel E Simmonds, Rebecca B Hoyle

## Abstract

Buruli ulcer (BU) is a neglected tropical skin disease caused by infection with *Mycobacterium ulcerans*. The mode of its transmission into the skin has not been fully clarified, but it is believed to involve mechanical transfer, perhaps by insect bites. *M. ulcerans* secretes a toxin called mycolactone which causes necrosis of the skin and formation of large skin lesions. Understanding the development of these skin lesions over time is crucial to understand the pathogenesis of BU. We formulate a mathematical model to describe the space-time dynamics of *M. ulcerans* bacteria, mycolactone and skin cell densities as well as fibrin deposition. We investigate the relative contributions to ulcer formation of direct cytotoxicity of mycolactone and ischaemia following the loss of coagulation control. Our results are consistent with existing biological evidence that higher mycolactone concentrations accelerate cell death. Furthermore, the model indicates that the rate of wound expansion is primarily controlled by the diffusion and net birth rate of bacteria, which sets the overall scaling for the speed of wound expansion, along with two dimensionless constants that define the shape of the trajectory of wound edge position against time, namely the ratio of the mycolactone diffusion coefficient to the bacterial diffusion coefficient, and the ratio of the mycolactone degradation rate to the bacterial net birth rate. When mycolactone diffuses significantly faster than bacteria spread or when mycolactone degrades more slowly than bacteria reproduce, the initial development of the wound is faster. In contrast, the long-term wound expansion speed appears to be relatively unchanged across a wide range of the dimensionless parameters.

**Author summary:** Buruli ulcer is a neglected tropical skin disease caused by *Mycobacterium ulcerans*, a bacterium that produces a toxin called mycolactone. The disease can cause extensive skin lesions, but the processes controlling how these lesions develop and expand are not fully understood. We developed a mathematical model to investigate how bacteria and mycolactone spread through the skin and contribute to skin-cell death. The model considers two possible mechanisms of tissue damage: direct toxic effects of mycolactone on skin cells and indirect damage caused by loss of blood supply following effects on endothelial cells. Our results show how differences in the sensitivity of skin-cell types to mycolactone can influence lesion development and suggest that indirect damage through ischaemia may contribute to tissue death even when mycolactone concentrations are too low to directly kill some cells. We also find that faster mycolactone diffusion or slower degradation can accelerate the initial development of a wound, while having relatively little effect on its longer-term expansion rate.

## Introduction

Buruli ulcer (BU) is a neglected tropical disease of the skin caused by *Mycobacterium ulcerans*, predominantly found in West and Central Africa, Australia and Japan [1, 2]. It is the third most common mycobacterial disease globally, but is significantly under-reported, especially in countries not reporting to the WHO. BU manifests initially as painless nodules, plaques, or oedema, progressing into ulcers. In Africa, the disease is associated with proximity to contaminated water sources, where *M. ulcerans* is linked to aquatic environments [3–5]. In Australia, possums appear to be the main environmental reservoir instead, and there is an association with mosquitoes instead of water. Recent evidence from southeastern Australia supports a role for mosquitoes in the transmission of *M. ulcerans* between possums and humans [6].

BU is characterized by necrosis of the skin’s dermis and subcutis, beginning with closed lesions and progressing to open ulcerative lesions with undermined skin edges [7]. Delays to diagnosis, which can be common in endemic rural areas, result in increased severity of cases. Treatment involves a 56-day regimen of rifampicin and clarithromycin, along with adaptive wound care, and surgery if necessary [8, 9]. The WHO recommends PCR confirmation before treatment, but accessibility in rural endemic areas remains challenging. Late-stage presentation complicates treatment and can result in long-term disability.

The characteristic pathology of BU lesions features coagulative necrosis and fibrin deposition distant from the infecting bacteria that are largely extracellular [10]. This is attributed to mycolactone, a diffusible exotoxin produced by *M. ulcerans* [11, 12] that plays a crucial role in host colonization and tissue damage. The predominant mechanism by which mycolactone drives BU is due to its ability to block the action of the essential cellular molecule Sec61 [13, 14]. This activity is immunosuppressive and cytotoxic, although mycolactone can also activate pro-inflammatory, Sec61-independent inflammasome signalling in macrophages, including through the IL-1*β* pathway [15, 16]. However, it remains unclear exactly how all the various consequences of Sec61 blockade by mycolactone identified *in vitro* result in the coagulative necrosis observed in *M. ulcerans*-infected skin. This is the problem addressed in the present study.

Skin tissue is complex and consists of multiple cell types with specialized functions [17]. The effect of mycolactone on many of these cells has been studied *in vitro* [18]. In the current work, for simplicity we have defined these as endothelial cells, which line the blood vessels and form capillaries, and stromal cells that represent all other cell types.

Endothelial cells are vitally important in tissue health. They regulate blood clotting within blood vessels in the absence of tissue damage [19], as well as controlling the permeability of vessels to blood and lymph components, as well as immune cells [20]. Previous work has shown that they are highly sensitive to as little as 2 ng ml^−1^ of mycolactone [21], which disrupts their normal function by inhibiting its anticoagulant functions and causing the destruction of the surrounding extracellular matrix leading to increased vascular permeability [22]. Endothelial cells are predominantly located in the dermis [17], the main site for bacterial growth in BU. It was recently suggested that the effect on endothelial cells may be linked to the fibrin deposition, since fibrin penetration of vessels is a feature of early lesion in the mouse model [22], and patient skin.

Other skin-resident cell types that have been tested for mycolactone-dependent cytotoxicity include keratinocytes, Schwann cells, fibroblasts and adipocytes [18]. They are all reported to be less sensitive to mycolactone than endothelial cells both *in vitro* and in patient skin [23]. These cell types found in the skin epidermis or dermis also do not directly influence blood coagulation control, but they collectively represent the majority of non-immune cells making up skin.

We consider that coagulative necrosis, observed in BU skin tissue, may result from two mycolactone-dependent mechanisms: (a) a direct mechanism due to cytotoxicity to exposed cells, and/or (b) an indirect mechanism due to tissue ischaemia. The two pathways are summarised in Figure 1, and will be described in detail below.

**Fig 1.**
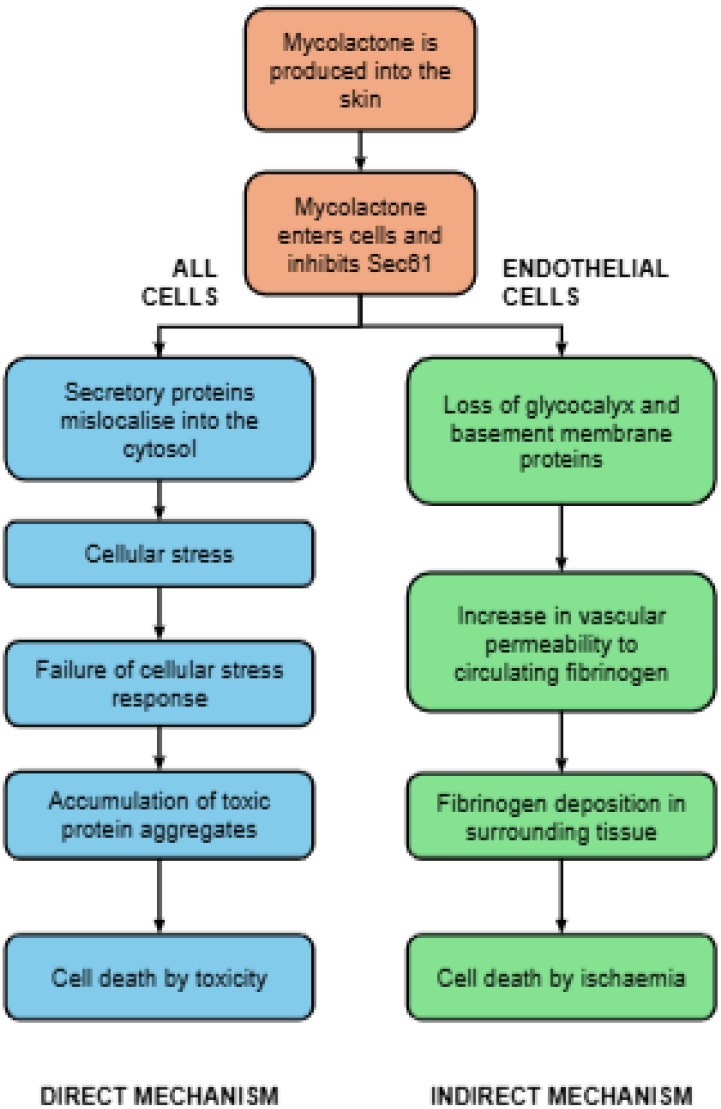
Cell death processes. A flow chart illustrating the direct and indirect mechanisms of mycolactone-dependent cell death processes in the skin exposed to mycolactone that are investigated in the current work. The mechanisms are described in detail in the text.

Direct cytotoxicity due to mycolactone is well described and arises from its action at the protein conducting channel Sec61. The underlying mechanism involves loss of membrane and secretory proteins and the cytosolic build-up of mislocalised proteins, due to disruption of protein transport into the endoplasmic reticulum [13]. The ubiquitin-proteasome system and autophagy, boosted by activation of cellular stress pathways, initially protects cells even as they loose adherence. However, eventually there is activation of caspase pathways leading to apoptotic cell death. [13, 24]. The amounts of mycolactone that cause direct cytotoxicity are reported to be different in endothelial cells (2 ng/ml) and stromal cells (125 ng/ml).

Tissue ischaemia due to the indirect consequences of mycolactone’s action on endothelial cells involves a working model whereby the Sec61-dependent loss of endothelial cell surface proteins and extracellular matrix causes endothelial dysfunction. This consequently allows plasma components (including the major precursor of fibrin clots, fibrinogen) into the stroma, resulting in fibrin deposition in tissue and causing cell death through ischaemia. Evidence from *in vitro* experiments, BU patient skin biopsies and *in vivo* models supports this working model. For instance, mycolactone causes the Sec61-dependent loss of the anticoagulant protein thrombomodulin, junction/adherens proteins as well as proteoglycans from the cell surface [21, 23].

Moreover, the loss of golgi-resident enzymes that manufacture the high order glycosylation of proteins causes rapid depletion of heparan sulfate and chondroitin sulfate. Together this results in the breakdown of the basement membrane *in vivo*, and an increase in vascular permeability [22]. Of particular interest is the penetration of fibrin into the deeper layers of larger blood vessels during early lesions, prior to widespread fibrin deposition at later stages.

In this manuscript, we present a mathematical model analyzing the formation of BU lesions, considering *M. ulcerans* growth dynamics, mycolactone production, and the potential contributions of direct cytotoxicity and ischaemia mechanisms to cell death.

Our study investigates these mycolactone-dependent mechanisms to unravel BU pathogenesis, understand the drivers of cell death, and offer insights into managing BU disease.

## Methods

### Model Formulation

Our model describes the formation of skin ulcers over time, assuming the skin comprises two cell types: endothelial cells (*n*_*e*_(*x, t*)) and stromal cells (*n*_*s*_(*x, t*)), where *x*∈ ℝ^2^ is a spatial coordinate in the plane of the skin, and *t* ≥0 is time. We make the simplifying assumption that on the scale of a BU wound, we can treat the distribution of skin cells as spatially mixed with a uniform ratio of the number of stromal cells to endothelial cells in healthy skin. We further assume that no new cells grow during ulceration due to the irreversible inhibitory effects of mycolactone on the cell cycle [13, 25, 26].

We denote bacterial density and local mycolactone concentration as *u*(*x, t*) and *m*(*x, t*), respectively. Both skin cell types are sensitive to mycolactone, with endothelial cells exhibiting higher sensitivity [13, 21]. When mycolactone concentration (*m*(*x, t*)) surpasses a threshold, skin cells undergo direct cytotoxicity-induced death over a 4-day period. The stromal cell toxicity threshold (*c*_*s*_) exceeds that of endothelial cells (*c*_*e*_).

Endothelial cells experience ischaemia-induced death 1 day after mycolactone reaches *c*_*e*_, while nearby stromal cells also undergo ischaemic death owing to loss of oxygen supply due to fibrin deposition. Mycolactone depletes thrombomodulin, junction proteins and proteoglycans from endothelial cells within a day *in vitro*. Although the precise rate of these events in tissue is unknown, we estimated the entire ischaemic cell death process to take 24 hours.

In our model we assume that direct contact of mycolactone with skin cells leads to cell death at constant rates *d*_*e*_ (for endothelial cells) and *d*_*s*_ (for stromal cells) if mycolactone concentration, *m*, is above the respective toxicity thresholds *c*_*e*_ and *c*_*s*_. In addition when mycolactone is above the endothelial cell toxicity threshold *c*_*e*_, fibrin accumulates causing cells to die through ischaemia at a constant rate *d*_*k*_.

The movement of bacteria and mycolactone through the skin are modelled by diffusion processes with diffusion coefficients *D*_*u*_ and *D*_*m*_ respectively. Bacterial cells are assumed to grow with a constant birth rate *λ*_*u*_ and die with a density-dependent death rate *K*_*u*_*u*^2^. Mycolactone is modelled as being produced by bacterial cells at a constant rate *λ*_*m*_ and decaying at a constant rate *K*_*m*_.

### Model equations

The resultant model equations for the dynamics of bacterial cells, mycolactone, endothelial cells and stromal cells are given by

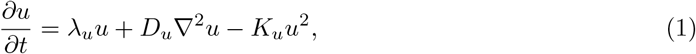

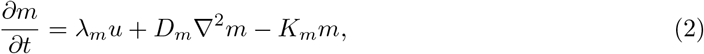

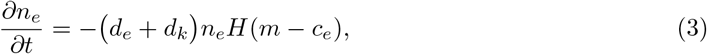

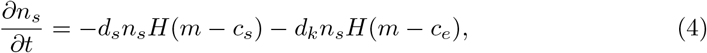

where *H*(*m* − *c*_*i*_) is the Heaviside step function that takes the value 0 for *m* ≤ *c*_*i*_ and 1 for *m > c*_*i*_. The variables and parameters used in the equations are listed in Table 1.

**Table 1.** Description of variables and parameters used in the model.

| Variable | Description |
| --- | --- |
| $n_e(x, t)$ | Endothelial cell density |
| $n_s(x, t)$ | Stromal cell density |
| $u(x, t)$ | Bacterial cell density |
| $m(x, t)$ | Mycolactone concentration |
| $x$ | Spatial variable (Distance) |
| $t$ | Time |
| Parameter | Description |
| $d_e$ | Death rate by direct cytotoxicity for endothelial cells |
| $c_e$ | Threshold value of mycolactone for cytotoxicity in endothelial cells |
| $d_k$ | Death rate by ischaemia for all cells |
| $d_s$ | Death rate by direct cytotoxicity for stromal cells |
| $c_s$ | Threshold value of mycolactone for cytotoxicity in stromal cells |
| $\lambda_u$ | Constant net birth rate by which bacteria are produced |
| $D_u$ | Diffusion coefficient of bacteria |
| $K_u$ | Constant for density-dependent bacterial death rate |
| $\lambda_m$ | Constant rate at which mycolactone is produced by bacterial cells |
| $D_m$ | Diffusion coefficient of mycolactone |
| $K_m$ | Degradation rate for mycolactone |
| $L$ | Typical length scale of ulceration |
| $\sigma$ | Standard deviation of the initial Gaussian distribution of bacteria |
| $\mu$ | Mean value of the initial Gaussian distribution of bacteria |

### Initial conditions

Skin cell densities are assumed to be initially uniform and non-zero such that *n*_*e*_(*x*, 0) = *n*_*e*0_, *n*_*s*_(*x*, 0) = *n*_*s*0_ and mycolactone concentration is initially zero, *m*(*x*, 0) = 0. We model the introduction of an initial colony of bacteria *u*(*x*, 0) = *u*_0_.

The endothelial cell density in healthy skin was estimated to be *n*_*e*0_ = 1.0 *×* 10^−3^ g.cm^−3^ and stromal cell density is 9 times that of density of endothelial cells (*n*_*s*0_ = 9*n*_*e*0_) [27], with the total cell density in healthy skin being 1.0 *×* 10^−2^ g.cm^−3^.

### Boundary conditions

We assume that the wound propagates into healthy skin where there are no bacteria or mycolactone. On an infinite domain of healthy skin the boundary conditions for *u* and *m* would be *u*(*t, x*) → 0 as *x*→*±*∞and *m*(*t, x*) → 0 as *x* →*±*∞. We will approximate these on a finite domain [0, *L*] by

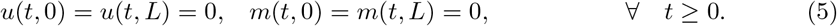

### Nondimensionalised model equations

The model equations can be non-dimensionalised by setting

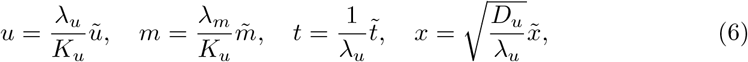

leading to

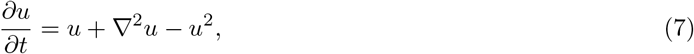

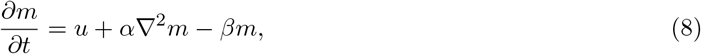

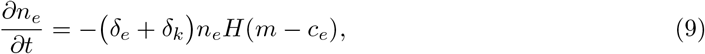

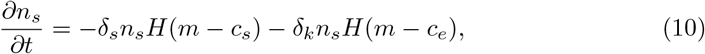

where the tildes have been dropped for convenience and where

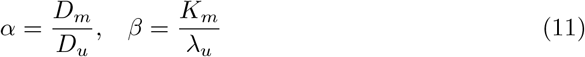

and

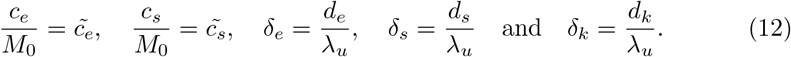

Initial and boundary conditions are nondimensionalised correspondingly. In nondimensional form the initial colony of bacteria is given by

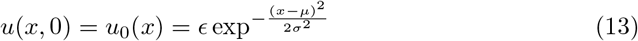

with *ϵ* = 1.0 *×* 10^−6^, *σ* = 1 and 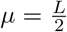 .

Note that Equation (7)) for the bacterial population is a form of the Fisher KPP equation [28], commonly used to describe the propagation of biological populations in space, which includes movement by diffusion and logistic growth. It has constant speed travelling wave solutions, which in our case represent the spreading of the bacteria across or through the skin.

Nondimensionalisation shows that, up to a scaling given by Equation (6), the dynamics of mycolactone propagation and hence of the position of the wound edge depends only on two non-dimensional parameters *α* and *β*.

### Model Parameterisation

We obtained certain parameter values from available experimental data and estimated others using similar observations from previous studies. The selection of specific parameter values is based on:

1. The doubling time of *M. ulcerans* is generally well-established as 2 days [29–31], corresponding to a growth rate of *λ*_*u*_ = 4.011 *×* 10^−6^s^−1^.
2. The rate at which mycolactone is produced by *M. ulcerans* (*λ*_*m*_) is not known. Therefore we estimated this based on experimental studies of mycolactone abundance in planktonic suspension cultures compared to dry cell weight [29]. From Equation (2), neglecting diffusion and degradation we have

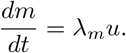

Focusing on the exponential growth phase of Figure 5A in [29] we let *m*_1_ and *m*_2_ be mycolactone concentrations at day 27 (*t*_1_) and day 39 (*t*_2_) and estimate values as 0.4 mgL^−1^ and 1.0 mgL^−1^ respectively. We estimated the value for *u* to be approximately between 10^−2^ and 0.5 gL^−1^ at these two time points. Using

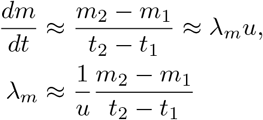

we have *λ*_*m*_ *≈* 1.157 *×*10^−9^s^−1^and *λ*_*m*_*≈* 5.787*×*10^−8^s^−1^ for the two values of *u*, and so for our model we choose as an order of magnitude estimate

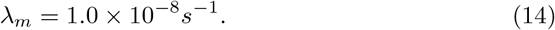
3. Assuming that 90% of the cells die through ischaemia within a day when the mycolactone concentration exceeds the endothelial cell toxicity threshold, corresponds to a cell death rate through ischaemia of *d*_*k*_ = 2.665 *×* 10^−5^s^−1^.
4. The rate constants *d*_*e*_, *d*_*s*_ and *d*_*k*_,corresponding to death by direct toxicity for endothelial and stromal cells and death by ischaemia respectively, are inversely proportional to the time for cell death. The time for cell death by direct toxicity is 4 days [21] compared to 1 day for death by ischaemia hence

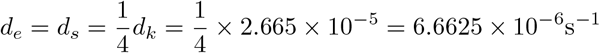
5. There is no experimental information available for diffusion rates of either mycolactone or *M. ulcerans* within infected skin. Indeed, as yet there is no experimental approach yet developed that can spatially determine the distribution in mycolactone in skin. Therefore diffusion rates were estimated based on the models of others, or work with other bacterial species in planktonic culture. These have estimated that the diffusion coefficient of a diffusible chemical is in the range 10^−9^ to 10^−6^ cm^2^ s^−1^ [32, 33]. We chose the diffusion coefficient of mycolactone to be *D*_*m*_ = 1.0*×* 10^−9^ cm^2^ s^−1^, reflecting its passive diffusion across cell membranes and uptake into ER membranes as observed by [34], and its irreversible binding to Sec61 [35].
6. To estimate the diffusion rate of bacteria in the skin, we use the dimensional scaling:

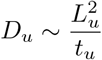

where *L*_*u*_ is a characteristic length scale and, *t*_*u*_ is a characteristic timescale for the diffusion. We estimate

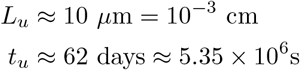

from Fig. 2B (inset) of [36]. Therefore we estimate

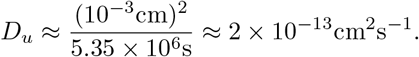

This calculation resulted in the value *D*_*u*_ *≈*2 *×*10^−13^cm^2^s^−1^. However, preliminary numerical simulations showed that this resulted in a propagation rate that was lower than expected, so we increased *D*_*u*_ until a more realistic propagation rate was achieved, using a final value of *D*_*u*_ = 1.87 *×*10^−11^ cm ^2^ s^−1^. Hence

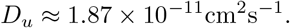
7. According to previous studies, the stromal cell toxicity threshold is 125 ng/ml [13, 24] which is equivalent to 1.25 *×* 10^−7^ g.ml^−1^ and the endothelial cell toxicity threshold is 4 ng/ml [21], which is equivalent to 4 *×* 10^−9^ g.ml^−1^.
8. We find the value of *K*_*m*_ using experimental work by [37] which suggests that

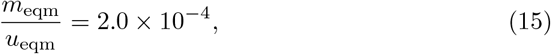

where *u*_eqm_ and *m*_eqm_ are the equilibrium values for bacterial density and mycolactone concentration in the spatially uniform case, respectively. From Eq. (2) we see that a spatially uniform equilibrium solution satisfies

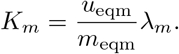

Substituting from Eq. (14) and Eq. (15) we obtain

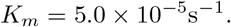
9. We choose *K*_*u*_ (the density-dependent bacterial death rate) by calibrating the model to produce equilibrium values of mycolactone concentration that illustrate the range of possible wound formation behaviours it permits. We acknowledge that this will not necessarily provide biologically reasonable values. To do this we first calculate the equilibrium bacterial density producing a given equilibrium mycolactone concentration from Eq. (15). Now choosing *m*_eqm_ = 1.5*×* 10^−7^gml^−1^ which is slightly above the sensitivity thresholds for the two cell types and will enable us to observe ulcer formation gives

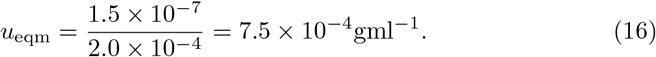

Then setting *u* = *u*_eqm_ in Eq. (1) gives

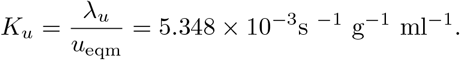

Similarly, choosing *m*_eqm_ = 6.0*×*10^−9^gml^−1^ which is a value slightly above the endothelial cell sensitivity threshold (but below the stromal cell sensitivity threshold) gives

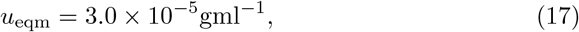

and then setting *u* = *u*_eqm_ in Eq. (1) gives

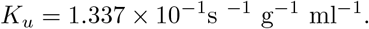

Thus we have two different values of *K*_*u*_ each corresponding to a different equilibrium mycolactone concentration. By choosing each of these values in turn we will be able to investigate how wound formation progresses when the mycolactone concentration surpasses the toxicity threshold for only endothelial cells, or when it is high enough to surpass the threshold for all cells.
10. According to [7], the length of a single small lesion in the WHO category one form of the BU disease can extend to 5 cm. Thus we model wounds growing up to 5 cm and stop the simulation when the wound reaches 5 cm. The simulation domain representing the skin into which the wound is growing is chosen to be larger than 5 cm so as to avoid boundary effects and allow the approximated boundary conditions Eq. (5) to be applied. However we show only the central 5cm portion of the domain containing the wound in our simulation results below.

### Values of the non-dimensional parameters

The non-dimensional parameters, *α* and *β* are calculated by substituting values of *D*_*m*_, *D*_*u*_, *K*_*m*_ and *λ*_*u*_ in Eqs. (11) giving

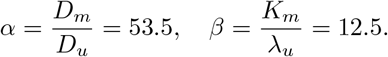

Values for *δ*_*e*_, *δ*_*s*_ and *δ*_*k*_ were obtained by substituting values of *d*_*e*_, *d*_*s*_ and *d*_*k*_ in Eq. (12) giving

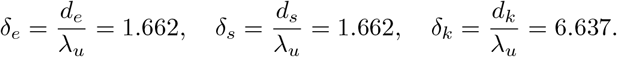

The non-dimensional value of *L* is given by

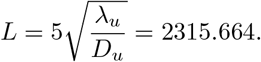

## Results

Equations (7), (8), (9), and (10) were solved numerically with initial and boundary conditions using Python’s “odeint” package in version 3.6. For Equations (7) and (8), we employed a central finite difference approximation for the second-order derivative in the spatial dimension, with *h* = *l/*(*N* −1). This led to a system of coupled ODEs, where *N* is the number of discretization points, and *l* is the domain length set to the typical ulceration scale *L*.

### Evolution of bacterial population and mycolactone concentration

We present plots for the solutions for bacterial (7) and mycolactone concentrations (8). The equations were solved in non-dimensional form and plotted in dimensional form.

In Figure 2, we start with a small amount of *M. ulcerans* bacteria which undergo logistic growth and diffusion, eventually reaching a plateau in the centre of the domain where the bacterial density remains approximately constant in time. We stop the simulation before the bacteria reach the edges of the wound.

When *M. ulcerans* bacteria grow, they produce mycolactone toxin which accumulates in the same pattern as the bacteria, due to the low diffusion rate. We set the mycolactone concentrations in the plateau or stationary phase *m*_eqm_ to be 6*×*10^−9^ gml^−1^ and 1.5 *×*10^−7^ gml^−1^. These values are slightly above the cell toxicity thresholds for the endothelial and stromal cells respectively. Mycolactone increases in the centre of the domain and diffuses across the domain with increase in time.

### Death of skin cells in response to mycolactone

#### Cell death by direct cytotoxicity only

We examine the effect of direct cytotoxicity on cell death by activating cell death through direct cytotoxicity and deactivating cell death by ischaemia in our model. Equations (9) and (10) then become:

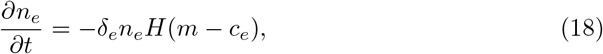

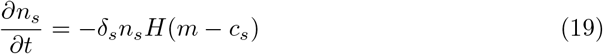

We substitute for parameter values and initial conditions, and Equation (18) and (19) are solved using the Python scipy “odeint” function to generate Figure 3.

**Fig 2.**
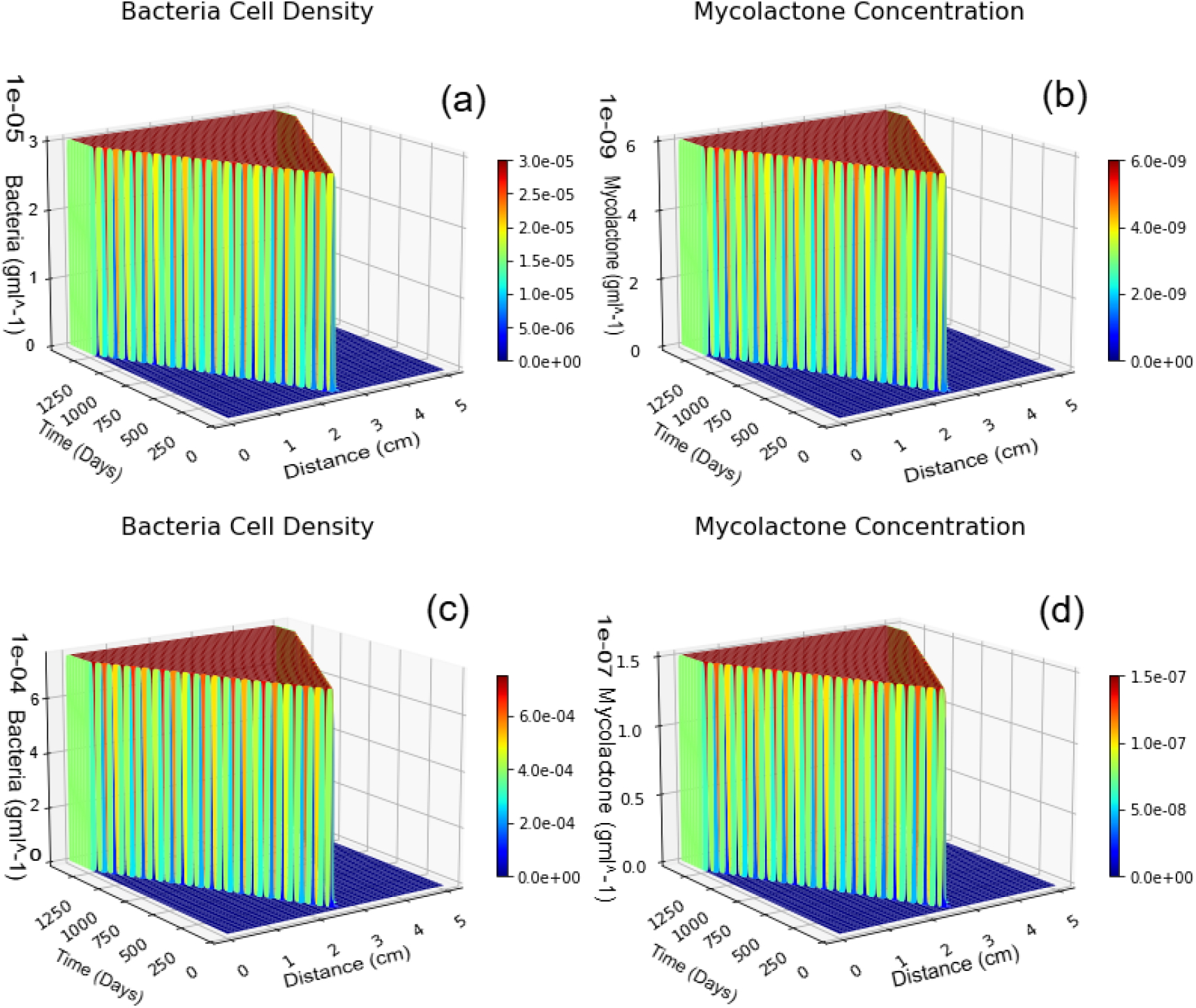
Evolution of bacterial population and mycolactone concentration in a given skin region over a period of 1250 days from an initial infection. In (a) and (b), the maximum bacterial population density at 3.0 *×* 10^−5^ g.cm^−3^ produces a mycolactone concentration of 6.0 *×* 10^−9^ g.cm^−3^. In (c) and (d), the maximum bacterial population density at 7.5 *×* 10^−4^ g.cm^−3^ produces a mycolactone concentration of 1.5 *×* 10^−7^ g.cm^−3^.

**Fig 3.**
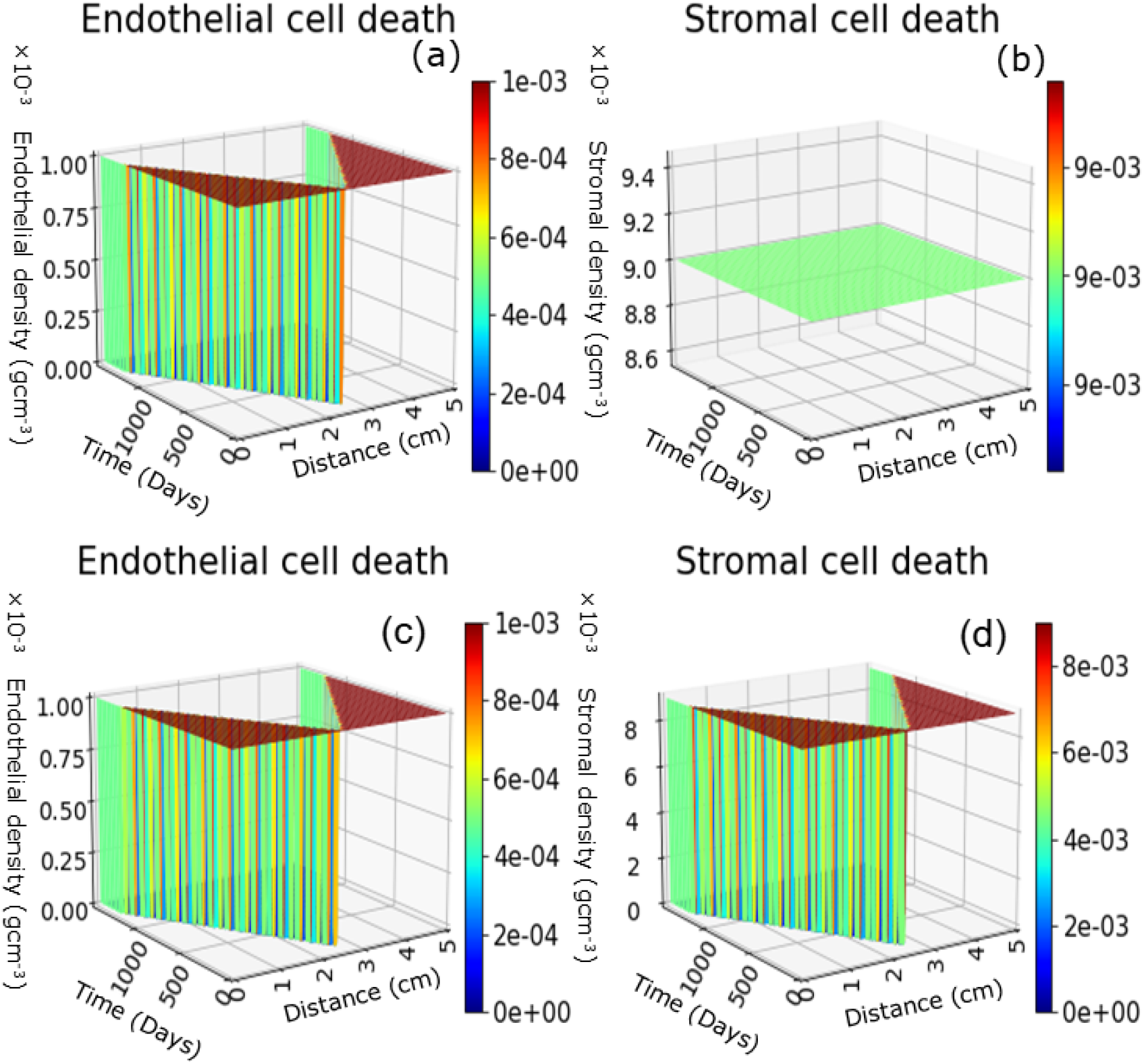
**Cell death by direct cytotoxicity:** In the first row, *m*_eqm_ = 6 *×* 10^−9^ gml^−1^, such that *c*_*s*_ *> m*_eqm_ *> c*_*e*_. In the second row, *m*_eqm_ = 1.5 *×* 10^−7^ gml^−1^, such that *m*_eqm_ *> c*_*s*_ *> c*_*e*_. We illustrate how (a) endothelial cells and (b) stromal cells respond to the presence of mycolactone.

In the first row of Figure 3, endothelial cells die, but no stromal cell death occurs because the mycolactone in the centre of the domain is above the toxicity threshold for endothelial cells, but below that for stromal cells that is (*c*_*e*_ *< m*_eqm_ *< c*_*s*_). In the second row we observe a fall in both endothelial and stromal cell densities in the centre of the domain due to the presence of mycolactone at a concentration above the toxicity threshold for both cell types (*m*_eqm_ *> c*_*s*_ *> c*_*e*_). The healthy skin regions observed at the left and right edges in Figure 3 occur because mycolactone has not yet reached the toxicity threshold in those regions.

The spatial variation in cell density arises because mycolactone values increase until they exceed the toxicity threshold in a given part of the domain and this process is controlled spatially by the diffusion of bacteria and the diffusion and secretion of mycolactone. Thus the wound develops gradually across the domain.

#### Cell death by ischaemia only

We examine the effect of ischaemia on cell death by activating cell death through ischaemia and deactivating cell death by direct cytotoxicity in the model. The equations for cell death in (9) and (10) become:

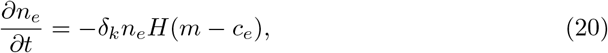

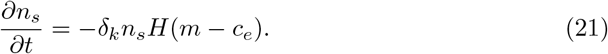

We substitute for parameter values and initial conditions, and Equation (20) and (21) are solved using the Python scipy “odeint” function to produce Figure 4.

**Fig 4.**
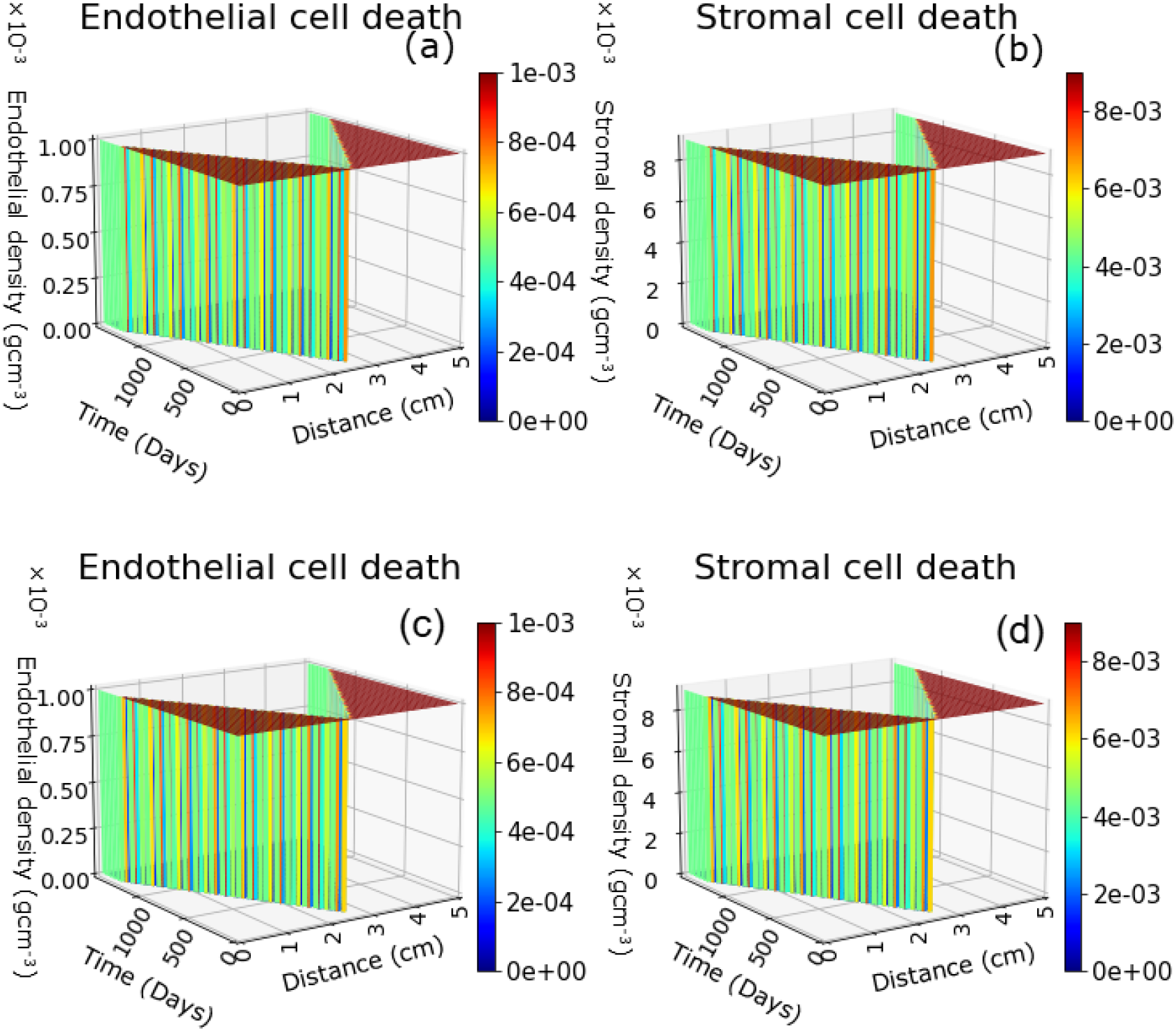
**Cell death by ischaemia:** In the first row, *m*_eqm_ = 6 *×* 10^−9^ gml^−1^, such that *c*_*s*_ *> m*_eqm_ *> c*_*e*_. In the second row, *m*_eqm_ = 1.5 *×* 10^−7^ gml^−1^, such that *m*_eqm_ *> c*_*s*_ *> c*_*e*_. We illustrate how (a) endothelial cells and (b) stromal cell respond tothe presence of mycolactone.

In Figure 4, we observe a significant and rapid reduction in skin cell densities in the centre of the domain in both sets of simulations because the mycolactone concentration is above the toxicity threshold for endothelial cells and this causes both cell types to die through ischaemia even though the direct toxicity threshold for the stromal cells has not been reached in the simulations shown in the first row. The regions of healthy skin observed at the left, and right edges are because mycolactone is not above the endothelial cell toxicity threshold there.

### Comparison between cell death by ischaemia and cell death by direct cytotoxicity for endothelial cells

To establish the difference between cell death by direct cytotoxicity (Figure 3), cell death by ischaemia (Figure 4) and cell death by both direct cytotoxicity and ischaemia, we visualise the comparison as indicated in Figure 5.

**Fig 5.**
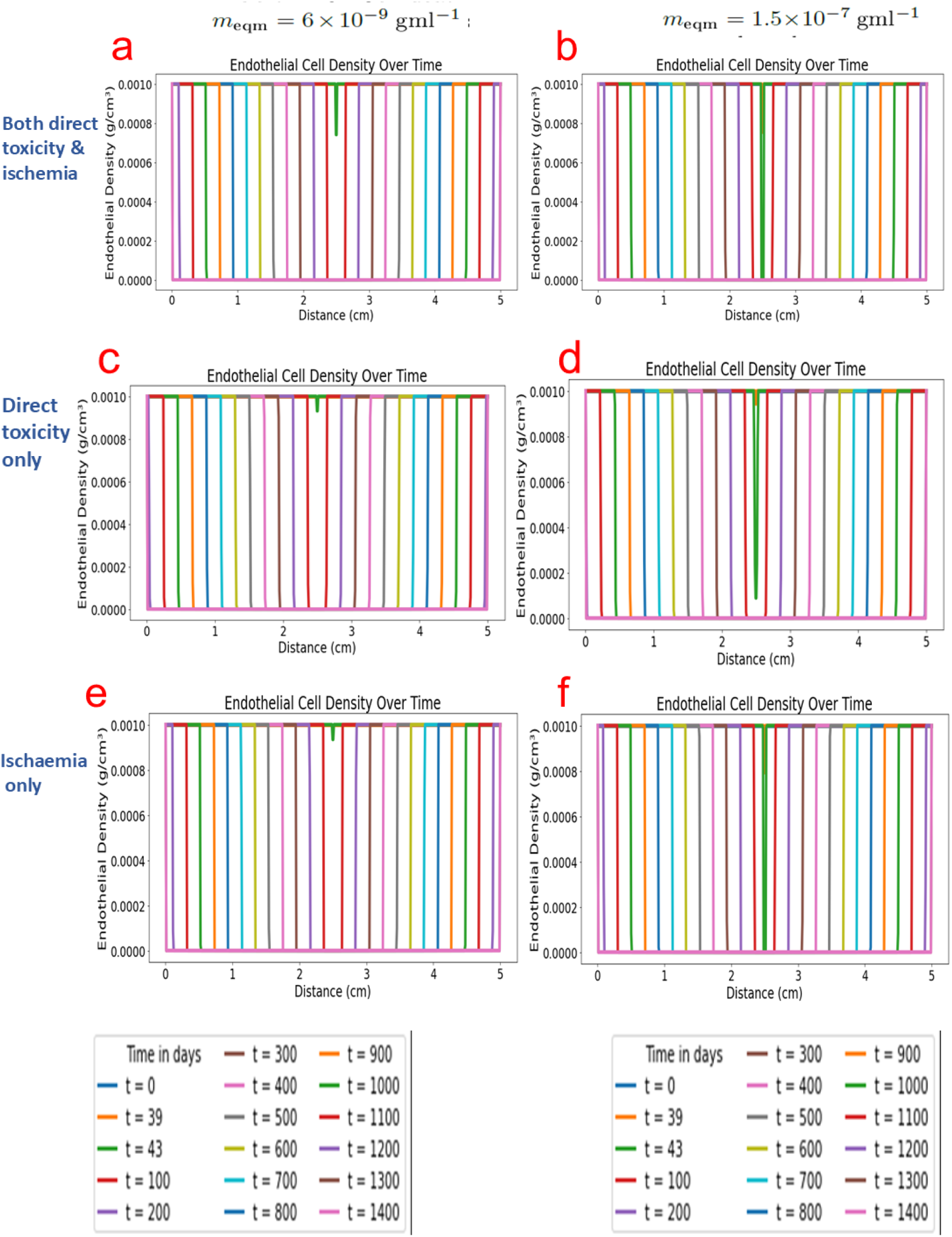
Comparison between cell death by ischaemia, direct cytotoxicity and both for endothelial cells. In the left column, *m*_eqm_ = 6.0 *×* 10^−9^ gml^−1^ and in the right column, *m*_eqm_ = 1.5 *×* 10^−7^ gml^−1^. This plot illustrates the variation in the endothelial cell density owing to exposure to mycolactone at chosen time points.

In comparing the three scenarios, our focus is on the time when cell density drops in the plots. We define the onset of cell-density decrease as the earliest time at which any point in the domain satisfies *n*(*x, t*) *< n*_0_. To determine this, we examine the spatial cell-density profile *n*(*x, t*) at each time point. For the case where *m*_eqm_ is set to 1.5*×*10^−7^ gml^−1^, the earliest drop in endothelial cell density in Figure 5 occurs at *t* = 39 days when both ischaemia and direct cytotoxicity mechanisms are included.

When *m*_eqm_ is set to 6 *×* 10^−9^ gml^−1^, the earliest drop happens at *t* = 43 days when both ischaemia and direct cytotoxicity operate simultaneously.

Our simulation reveals that cell death occurs earliest when *m*_eqm_ is set to 1.5 *×* 10^−7^ gml^−1^ with both direct cytotoxicity and ischaemia. The latest occurrence is when *m*_eqm_ is set to 6.0*×* 10^−9^ gml^−1^ with direct cytotoxicity only.

The reason for earlier cell death for *m*_eqm_ = 1.5*×* 10^−7^ is that mycolactone surpasses the toxicity threshold more quickly compared to when *m*_eqm_ = 6 *×*10^−9^ gml^−1^.

Our results highlight how changes in mycolactone values relative to skin cell toxicity thresholds can influence the process of cell death observed in BU. These findings affirm existing knowledge on how higher mycolactone concentrations accelerate cell death. It also supports the hypothesis that endothelial cells may play a critical role in tissue necrosis due to their high sensitivity to mycolactone and specialist function within the skin.

### Speed of the wound lesion

We monitor the position of the wound edge over time. We define the position of the wound edge to be the location of a sudden drop in cell density. This occurs when mycolactone surpasses the relevant toxicity threshold, and marks the initiation of the wound at the center followed by its extension outward in both directions. The diminishing distance between the wound edges and the domain edges over time indicates an increase in wound size. The speed of the wound is obtained by evaluating the slope of the wound edge position curve:

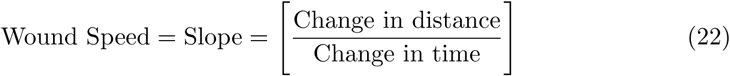

Equation 22 represents the ratio of how far the wound edge has moved to the amount of time it took for that movement to occur. It provides a basic representation of the rate at which the wound is spreading over time.

Figure 6 illustrates the progression of the wound front as the density of the skin cells decreases. The slope of this curve in this representation quantifies the wound speed, indicating the rate at which the wound forms.

**Fig 6.**
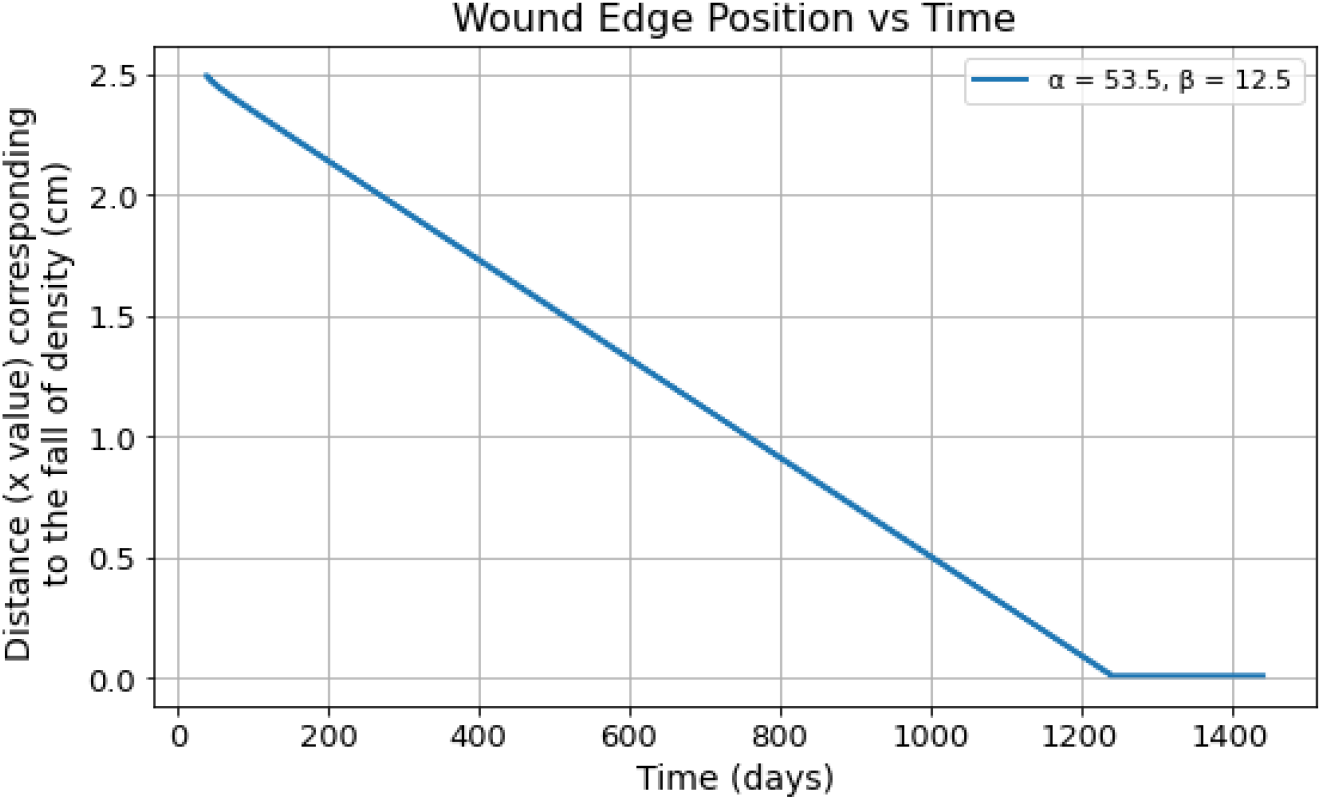
**Visualization of wound progression over time:** Starting from the center of the domain (*x* = 2.5 cm), we track the leftward movement of the wound edge over time. The wound edge is defined as the first spatial location where the endothelial cell density drops below its initial value. At each time step, this position is recorded, producing a set of distance–time data points. Using the parameter values (*α, β* = 53.5, 12.5) with spatial and temporal resolutions *N* = 500 and *N*_*t*_ = 30000 respectively. We focus on the approximately linear region of the wound progression between *t* = 600 days and *t* = 1000 days. The wound speed is then calculated as the slope of the distance-time curve over this interval, using Equation (22), yielding a value of −2.05*×*10^−3^ cm/day. The negative sign indicates that the wound edge is moving leftwards (towards decreasing *x*). The remaining parameter values are as given in Table 2.

#### Time scaling and interpretation

Time in the model is nondimensionalised using the characteristic timescale ^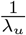^, where *λ*_*u*_ represents the constant net birth rate by which bacteria are produced. Dimensional time is recovered via 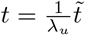, so that small values of *λ*_*u*_ lead to large physical times even for only moderately long dimensionless simulations. This scaling reflects the assumption that bacterial proliferation governs the dominant dynamics of the system.

**Table 2.** Parameter values used in the model.

| Notation | Dimensional value | Source |
| --- | --- | --- |
| $\lambda_u$ | $4.011 \times 10^{-6} \text{ s}^{-1}$ | Estimated in item 1 |
| $\lambda_m$ | $1.0 \times 10^{-8} \text{ s}^{-1}$ | Estimated in item 2 |
| $d_k$ | $2.665 \times 10^{-5} \text{ s}^{-1}$ | Estimated in item 3 |
| $d_s$ | $6.6625 \times 10^{-6} \text{ s}^{-1}$ | Estimated in item 4 |
| $d_e$ | $6.6625 \times 10^{-6} \text{ s}^{-1}$ | Estimated in item 4 |
| $D_m$ | $1.0 \times 10^{-9} \text{ cm}^2 \text{ s}^{-1}$ | Estimated in item 5 |
| $D_u$ | $1.87 \times 10^{-11} \text{ cm}^2 \text{ s}^{-1}$ | Estimated in item 6 |
| $K_m$ | $5.0 \times 10^{-5} \text{ s}^{-1}$ | Estimated in item 8 |
| $K_u$ | $5.348 \times 10^{-3} \text{ s}^{-1} \text{ g.ml}$ | Estimated in item 9 |
| $K_u$ | $1.337 \times 10^{-1} \text{ s}^{-1} \text{ g.ml}$ | Estimated in item 9 |
| $L$ | 5 cm | Estimated in item 10 |
| $c_e$ | $4.0 \times 10^{-9} \text{ g.ml}^{-1}$ | From [21] |
| $c_s$ | $1.25 \times 10^{-7} \text{ g.ml}^{-1}$ | From [13, 21, 24] |

**Table 3.** Non-dimensional parameters used in the model.

| Notation | Description | Value |
| --- | --- | --- |
| $\alpha$ | Ratio of the mycolactone to bacterial diffusion coefficients | 53.5 |
| $\beta$ | Ratio of the mycolactone decay rate to the bacterial growth rate | 12.5 |
| $\delta_e$ | Ratio of death rate (endothelial cell) to birth rate by which bacteria is produced | 1.662 |
| $\delta_s$ | Ratio of death rate (stromal cell) to birth rate by which bacteria is produced | 1.662 |
| $\delta_k$ | Ratio of death rate by ischaemia to constant net birth rate by which bacteria are produced | 6.637 |
| $L$ | Length of the wound | 2315.664 |

In regimes where the bacterial diffusion coefficient *D*_*u*_ is small, spatial propagation is slow and long simulation times are required to observe meaningful changes in bacterial density and toxin accumulation. The resulting extended timescales therefore arise naturally from the model structure and parameter choices, and are used here to examine qualitative disease progression rather than to make precise chronological predictions.

### The effect of non-dimensional parameters on the speed of wound propagation

We explore the sensitivity of model outputs to variations in the non-dimensional parameters *α* and *β*. Here *α* denotes the ratio of the diffusion coefficient of mycolactone (*D*_*m*_) to that of bacteria (*D*_*u*_) and *β* represents the ratio of the degradation rate of mycolactone (*K*_*m*_) to the bacterial growth rate (*λ*_*u*_). Hence, values of *α* greater than 1 represent scenarios where mycolactone is diffusing faster than the bacteria. Values of *β* greater than 1 represent scenarios where mycolactone is degrading faster than the bacteria grow.

For various *α* and *β* combinations, we track the position of the wound edge, calculating the slope of the resulting curve to determine the speed of wound progression. The simulation is conducted over a period of 1500 days. Simulations were performed with spatial resolution (*N* = 500) and temporal resolution(*N*_*t*_ = 30000), with the higher temporal resolution used to improve the accuracy of the numerical computations. The dimensionless parameters *α* and *β* were varied by changing *D*_*m*_ and *K*_*m*_; all other parameter values are given in Table 1.

The steady-state speed of the wound was calculated as the slope of the wound edge position versus time. For each simulation, we identified the time interval over which the wound edge position varied linearly in time, corresponding to an interval of approximately constant speed. This interval was chosen to exclude both the initial transient dynamics and the later stage where the wound edge approaches the domain boundary and stops moving. The start and end time points of the interval were determined individually for each parameter set based on where the curve exhibited approximately constant slope and we chose the interval between *t* = 600 days and *t* = 1000 days.

Figure 7 shows the wound-edge position over time when *α* is varied while *β* = 12.5 is held constant. Curves corresponding to very small values of *α* (e.g., 0.00535 and 0.0535) are omitted, as they lie indistinguishably on top of the curve for *α* = 0.535. For *α*≤ 5.35, the wound edge trajectories overlap almost exactly. This indicates that variations in *α* within this range have a negligible effect on wound dynamics.

**Fig 7.**
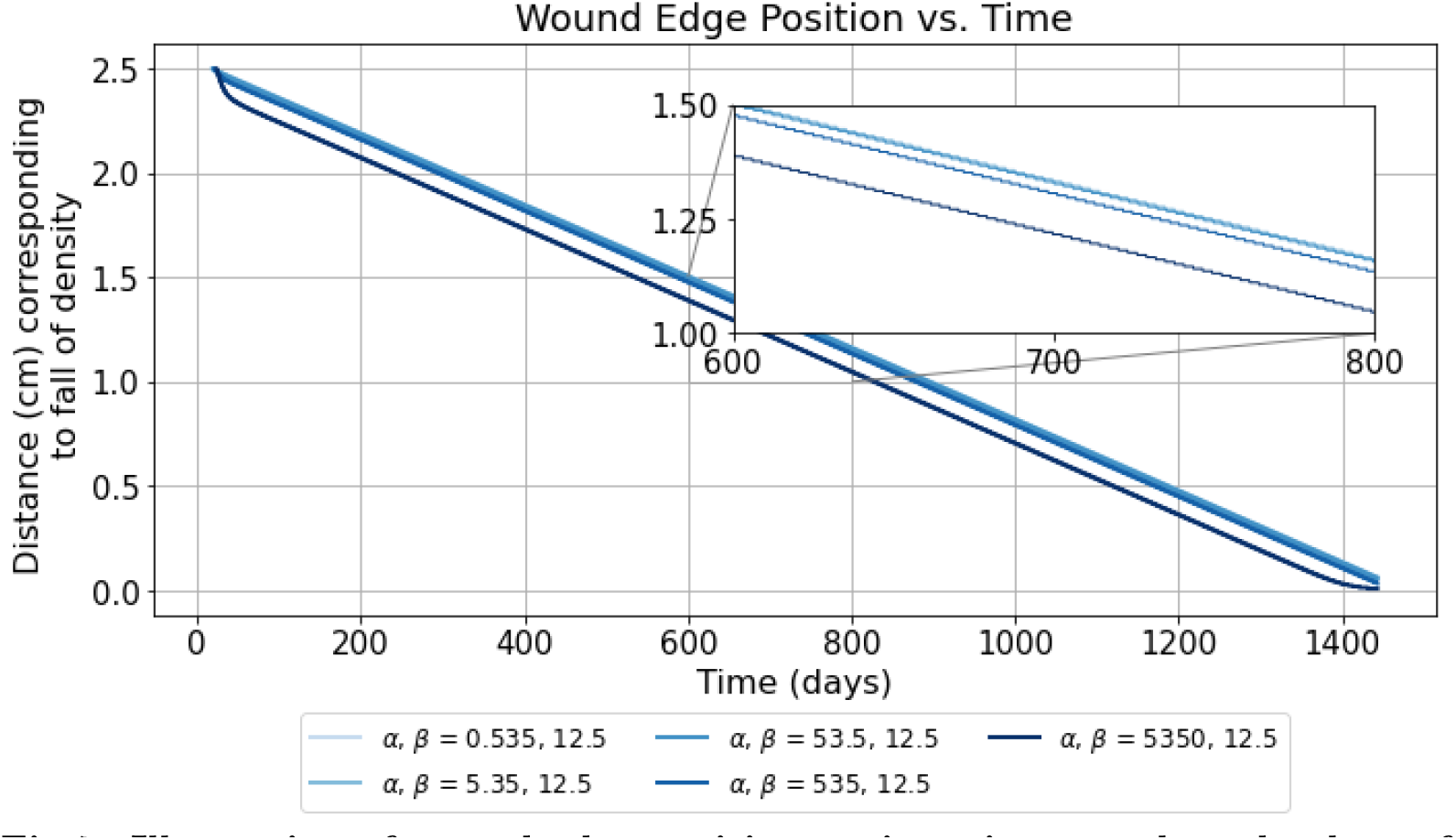
Illustration of wound edge position against time at selected values of the non-dimensional parameters *α* and *β*, where *β* = 12.5 is fixed and *α* varies.

The slopes corresponding to selected (*α, β*) combinations, shown in Table 4, represent the rate of wound edge progression over time. As *α* increases the slopes remain very close to −2.05*×* 10^−3^ cm/day to 3 sig. fig., indicating that increasing *α* has very little effect on the steady-state rate of wound expansion.

**Table 4.** Slopes for different values of *α* with *β* = 12.5.

| $\alpha$ | $\beta$ | Slope (cm/day)<br>to 3 sig. fig. |
| --- | --- | --- |
| 0.535 | 12.5 | $-2.05 \times 10^{-3}$ |
| 5.35 | 12.5 | $-2.05 \times 10^{-3}$ |
| 53.5 | 12.5 | $-2.05 \times 10^{-3}$ |
| 535 | 12.5 | $-2.02 \times 10^{-3}$ |
| 5350 | 12.5 | $-2.02 \times 10^{-3}$ |

However, there is an effect of *α* on the initial speed of wound expansion, with larger *α* leading to faster initial wound speed. In particular when *α* = 5350, the initial rate of expansion is noticeably faster than for smaller values of *α* (Figure 7).

These results suggest that mycolactone diffusion only significantly influences wound dynamics when it is much faster than bacterial diffusion. In this case, mycolactone can spread ahead of the bacteria, accelerating tissue damage and driving rapid initial wound expansion in regions the bacteria have not yet reached. As a result, the wound edge initially advances more rapidly, producing a steeper initial decline in the wound-edge curves. By contrast, when mycolactone diffuses at a rate comparable to or slower than bacterial diffusion, it remains more localised to the bacterial population, and wound propagation is instead dominated by bacterial movement. In this case, mycolactone either diffuses more slowly than the bacteria (*α <* 1) or faster, but to a lesser extent than for the largest values of *α*. It therefore remains concentrated nearer to the bacterial population, and the wound cannot advance beyond where the bacteria are located, leading to curves that overlap almost perfectly.

Figure 8 shows the wound-edge position over time for different values of β, where β is the ratio of the mycolactone decay rate to the bacterial growth rate.

**Fig 8.**
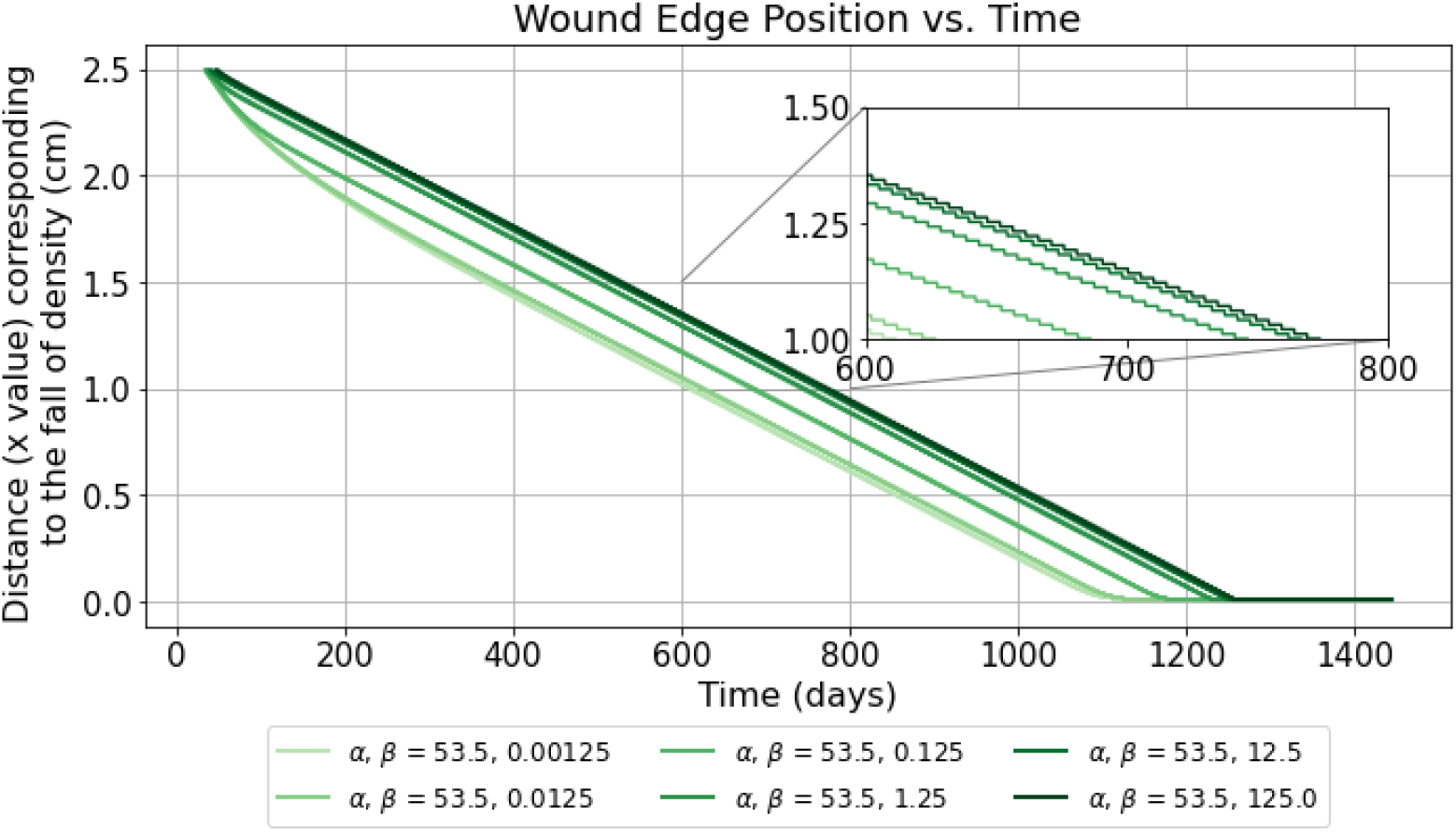
Illustration of wound edge position against time at selected values of the non-dimensional parameters *α* and *β*, where *α* = 53.5 is fixed and *β* varies.

The steady-state wound speed is very similar across all values of *β* (Table 5), indicating that *β* has little influence on the long-term wound expansion rate. However, the initial development of the wound is faster for smaller *β*. This is because smaller *β* corresponds to faster production of bacteria (and hence mycolactone) compared to degradation of mycolactone and so the concentration of mycolactone is expected to be transiently higher, leading to faster accumulation of tissue damage. Increasing *β* does not significantly change the wound expansion speed once progression is underway, but slows the initial wound development.

**Table 5.** Slopes for different values of *β* with *α* = 53.5.

| $\alpha$ | $\beta$ | Slope (cm/day)<br>to 3 sig. fig. |
| --- | --- | --- |
| 53.5 | 0.00125 | $-2.05 \times 10^{-3}$ |
| 53.5 | 0.0125 | $-2.05 \times 10^{-3}$ |
| 53.5 | 0.125 | $-2.03 \times 10^{-3}$ |
| 53.5 | 1.25 | $-2.03 \times 10^{-3}$ |
| 53.5 | 12.5 | $-2.05 \times 10^{-3}$ |
| 53.5 | 125.0 | $-2.05 \times 10^{-3}$ |

## Discussion

In this study, we developed a mathematical model to shed light on the dynamics of wound growth during *M. ulcerans* infection. We formulated model equations, defined variables, parameters, and their respective initial and boundary conditions, and we provided a comprehensive summary of our model. Utilizing numerical solutions, we illustrated the progression of bacterial growth, mycolactone concentration, and skin cell density, highlighting the spatial spread of lesions through a traveling wave solution.

Our findings illustrate the pivotal role that mycolactone plays in BU lesion formation. The simulations depict an exponential increase in the concentrations of *M. ulcerans* bacteria and mycolactone. By controlling the equilibrium concentration of mycolactone, we illustrate its differential effect on the direct cytotoxic death of different cell types. Hence, when the equilibrium concentration of mycolactone exceed the cytotoxicity threshold for both endothelial and stromal cells, both cell types die. On the other hand, when it exceeds only the cytotoxicity threshold of the endothelial cells, the stromal cells survive in the absence of ischaemic death. The differential sensitivity of mammalian cells with different specialised functions following exposure to mycolactone *in vitro* [18] is not simply a function of different laboratories and variations in experimental methodology. A recent side-by-side comparison of mycolactone cytotoxicity to cells *in vitro* recently demonstrated this range in detail [38]. The molecular explanation for this difference currently remains elusive.

On the other hand, in the postulated ischaemia-dependent indirect mechanism the high sensitivity of endothelial cells means that stromal cells can still die even when concentrations of mycolactone remain lower than that that would normally kill them. This suggests that this mechanism may be particularly important at driving tissue necrosis at the margins of the lesion where mycolactone concentrations are at their lowest.

A key outcome of nondimensionalisation of the model equations is that, aside from an overall scaling that depends on the diffusion coefficient and net birth rate of bacteria, wound propagation depends primarily on the dimensionless parameters *α* and *β*, rather than on properties of the mycolactone production and diffusion individually. Here, *α* represents the ratio of mycolactone to bacterial diffusion, while *β* captures the ratio of mycolactone decay to bacterial growth. This shows that transport processes along with mycolactone production and decay, influence lesion spread.

Sensitivity analysis shows that both *α* and *β* have limited influence on steady-state wound speed. However, the initial development of the wound is faster when *α* is high or *β* is low. For large *α*, mycolactone diffuses significantly ahead of the bacterial population, leading to faster initial wound expansion. This suggests that an increase in toxin mobility can accelerate tissue damage by extending cytotoxic effects into previously unaffected regions. Smaller values of *β*, correspond to slower mycolactone decay relative to bacterial growth, resulting in transiently higher mycolactone concentrations that accelerate initial wound expansion.

A significant limitation of our model is the general lack of experimental data that could be used support its parameterisation. For instance, while there is some data regarding mycolactone yields from planktonic cultures of *M. ulcerans*, an actual production rate is lacking. It is also unknown to what extent the rates of mycolactone production *in vitro* are replicated *in vivo*, which may change depending on the stage of infection (intramacrophage, or extracellular). Likewise, the issue of the stability of mycolactone within tissue is also an open question. Animals are able to tolerate injection of amounts of mycolactone that would be rapidly toxic to cells in culture, and the cellular effects *in vivo* seem more transient than they are *in vitro*.

Measuring mycolactone remains a major challenge in the field, due to its composition (a lipid-like polyketide lactone). Indeed, the field still awaits a methodological advance that can easily quantify mycolactone without first performing a lipid extraction on tissue, or detect it directly in skin to understand its spatial distribution. This would be required to fully understand the diffusion rate of mycolactone within skin tissue.

Another limitation is that we assumed that mycolactone and *M. ulcerans* would both reach equilibrium in skin, which may not hold in natural infections. We chose to do this so as to illustrate the differential impact of mycolactone on epithelial and stromal cells. However it would also be possible to examine the model under non-equilibrium conditions corresponding to the initial stages of bacterial growth and invasion. Furthermore, our model did not include any elements of the immune system, such as macrophage cells and neutrophils that may be important at early stages of infection, by phagocytosing and killing *M. ulcerans*. Therefore future research should focus on solving these issues, refining parameter values, and exploring model adjustments to reflect real-world conditions. The model could also be developed further in future to consider two horizontal dimensions laterally and a vertical dimension through the depth of the skin, along with more realistic spatial structure of the skin itself in terms of the distribution of skin cells. This would provide further insight into the details of wound formation.

The application of the model lies in enhancing understanding of BU lesion development and aiding predictions for tissue degradation. The study provides insights for future research into the control and management of BU, and contributes to the understanding of this neglected tropical disease.

## Acknowledgments

FA was supported by a University of Southampton Centrally Awarded Vice Chancellor’s scholarships. The funder had no role in study design, data collection and analysis, decision to publish, or preparation of the manuscript.

## AI Disclosure

ChatGPT (OpenAI) was used to assist with language editing and code documentation. All scientific content, analyses, and conclusions were developed, reviewed, and verified by the authors.

